# The RopGEF KARAPPO is Essential for the Initiation of Vegetative Reproduction in *Marchantia*

**DOI:** 10.1101/385682

**Authors:** Takuma Hiwatashi, Koh Li Quan, Yukiko Yasui, Hideyuki Takami, Masataka Kajikawa, Hiroyuki Kirita, Mayuko Sato, Mayumi Wakazaki, Katsushi Yamaguchi, Shuji Shigenobu, Hidehiro Fukaki, Tetsuro Mimura, Katsuyuki T. Yamato, Kiminori Toyooka, Shinichiro Sawa, Daisuke Urano, Takayuki Kohchi, Kimitsune Ishizaki

## Abstract

Many plants can reproduce vegetatively, producing clonal progeny from vegetative cells; however, little is known about the molecular mechanisms underlying this process. Liverwort (*Marchantia polymorpha*), a basal land plant, propagates asexually via gemmae, which are clonal plantlets formed in gemma cups on the dorsal side of the vegetative thallus [1]. The initial stage of gemma development involves elongation and asymmetric divisions of a specific type of epidermal cell, called a gemma initial, which forms on the floor of the gemma cup [2, 3]. To investigate the regulatory mechanism underlying gemma development, we focused on two allelic mutants in which no gemma initial formed; these mutants were named *karappo*, meaning “empty”. We used whole-genome sequencing of both mutants, and molecular genetic analyses to identify the causal gene, *KARAPPO* (*KAR*), which encodes a Rop guanine nucleotide exchange factor (RopGEF) carrying a PRONE catalytic domain. *In vitro* GEF assays showed that the full-length KAR protein and the PRONE domain have significant GEF activity toward MpRop, the only Rop GTPase in *M. polymorpha*. Moreover, genetic complementation experiments showed a significant role for the N- and C-terminal variable regions in gemma development. Our investigation demonstrated an essential role for KAR/RopGEF in the initiation of plantlet development from a differentiated cell, which may involve cell polarity formation and subsequent asymmetric cell division via activation of Rop signaling, implying a similar developmental mechanism in vegetative reproduction of various land plants.

## Results and Discussion

### Gemma development in *Marchantia polymorpha*

Vegetative reproduction is a form of asexual reproduction in which clonal individuals develop directly from vegetative tissues, such as leaves, stems, and roots. Vegetative reproduction is a developmental process based on totipotency, which is the potential for a cell, even a differentiated cell, to regenerate organs or whole plantlets [4-6]. Many plants in diverse lineages exhibit vegetative reproduction, *e.g.* potato (*Solanum tuberosum*), which produces tubers in underground stems, *Kalanchoe diagremontiana*, which forms plantlets at the leaf margins, the Dhalia family, which develop root tubers, and the hen and chicken fern (*Asplennium bulbiferum*), which grows small bulbils on the top of fronds [7]. However, very little is known about the underlying molecular mechanisms of vegetative reproduction.

One of the most basal lineages in extant land plants, the liverwort *Marchantia polymorpha*, has the ability to propagate asexually by forming clonal plantlets, called gemmae, in a cupule or “gemma cup”, a cup-like receptacle formed on the dorsal side of the thallus, which is the gametophyte plant body (Figure S1A). The development of the gemma and gemma cup in *M. polymorpha* has been described on the basis of histological observations [2, 8]. In the basal floor of the gemma cup, epidermal cells undergo cell elongation followed by two cycles of asymmetrical cell division to form an apical gemma cell and a basal cell (Figure S1E). The gemma cell continues to divide and finally produces the discoid gemma with two laterally developed apical notches. The basal cell does not divide any further and differentiates into a stalk cell [1, 2]. Mucilage papillae also develop from individual epidermal cells located in the floor of gemma cups [2, 3]. In our histological observations, various stages of developing gemmae and single-celled mucilage papillae (large club-shaped cells) were observed in the basal floor of the gemma cup (Figure S1B–D). The elongated morphology of mucilage papillae was distinct, with a number of single-membrane vesicles in their cytosol (Figure S1C). At the initial stage of gemma development, the basal stalk cell was already vacuolated, and the apical gemma cell underwent several rounds of periclinal cell divisions. In most cases, an anticlinal cell division was observed in the basal floor cell attached to the early stage of the developing gemma, while there was no cell division observed in the basal floor cell attached to the mucilage cell (Figure S1B–E).

### Isolation of *karappo-1* and *karappo-2* mutants

In recent years, *M. polymorpha* has been exploited as a basal plant model system due to the availability of whole-genome sequence information, high-efficiency transformation methods, and genetic modification techniques [9-16].

To identify key regulator(s) involved in the initial stage of gemma development in *M. polymorpha*, we focused on two mutants, named *karappo-1* (*kar-1*) and *karappo-2* (*kar-2*), that show a common phenotype of impaired gemma formation. These two mutants were isolated independently; *kar-1* was isolated during the screening of T-DNA-tagged lines for morphological phenotypes of the gametophyte thallus [17], and *kar-2* was isolated from transgenic lines generated by biolistic delivery of a plasmid [18]. In the wild type as well as in *kar-1* and *kar-2* mutants, gemma cups formed at intervals on the dorsal side of thalli along the midrib (Figure 1A, B, and C). Numerous mature gemmae were observed from the top of the wild-type gemma cup; however, no gemmae were found in the *kar-1* and *kar-2* mutants (Figure 1F, G, and H). Transverse sections of the gemma cup showed no developing gemmae in the gemma-cup of the *kar-1* and *kar-2* mutants (Figure 1L, M, Q, and R), while various stages of developing gemmae were observed in the wild type (Figure 1K and P). In contrast, mucilage papillae were formed from the basal epidermis of the gemma cup in the *kar-1* and *kar-2* mutant as in the wild type (Figure 1U, V, and W). There was no distinct impairment in the other aspects of vegetative development in the *kar-1* and *kar-2* mutants compared to the wild type (*i.e.* growth rate of thalli, air chamber formation, and rhizoid development). These observations suggest that the initial stage of gemma development is defective in *kar-1* and *kar-2* mutants.

**Figure 1.**
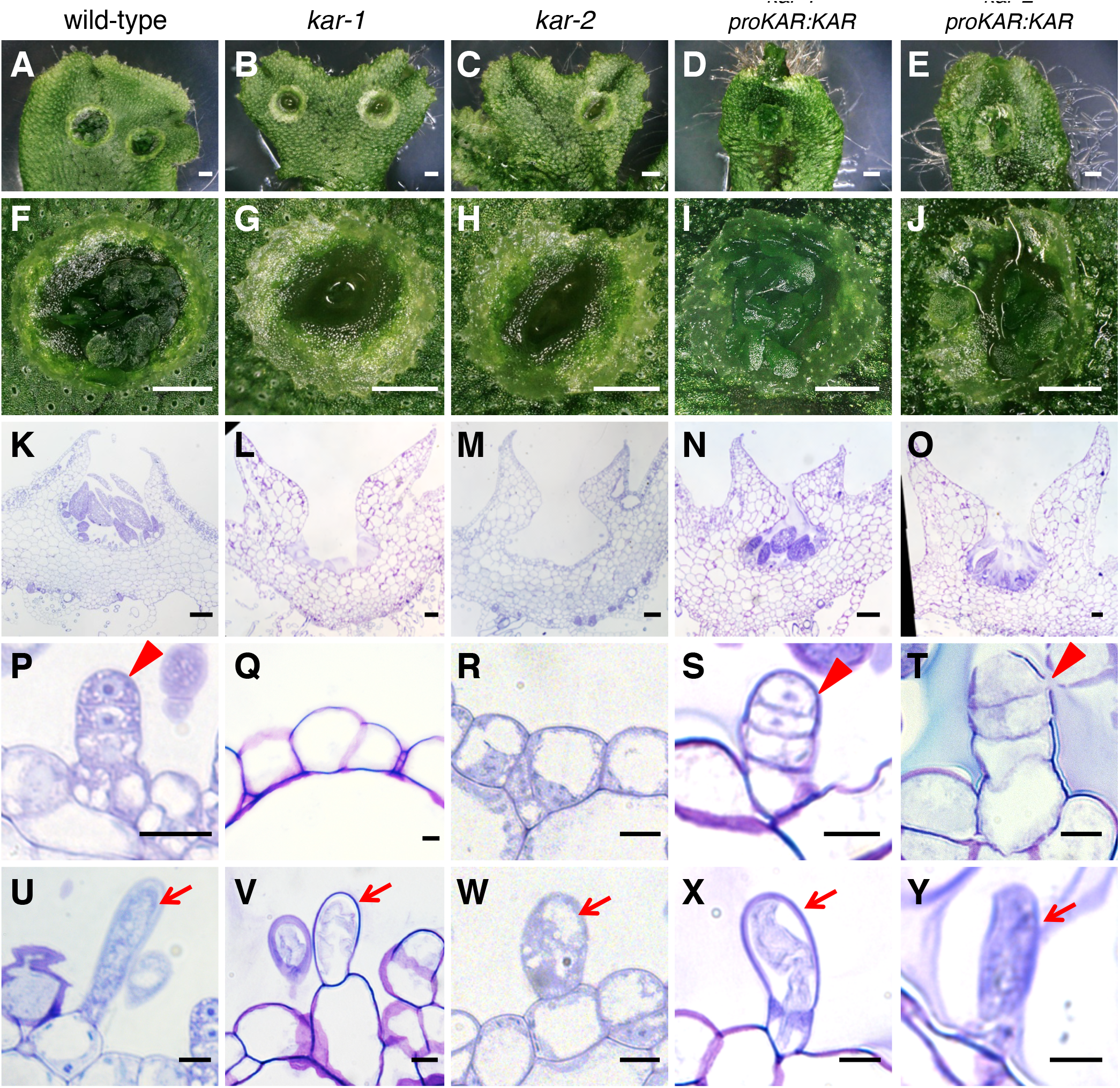
**Phenotype of the *kar* mutants and their complimented lines (A–Y)** Five genotypes are presented, one in each column: (first column) wild type, (second column) *kar-1*, (third column) *kar-2*, (fourth column) *kar-1* complementation line (*kar-1* transformed with *proKAR:KAR*), (fifth column) *kar-2* complementation line (*kar-2* transformed with *proKAR:KAR*). **(A–E)** Top view of 2-week-old thalli grown from tips of thalli. Scale bars represent 1 mm. **(F–J)** Surface view of gemma cups in 2-week-old thalli. Scale bars represent 1 mm. **(K–O)** Toluidine-blue-stained transverse sections of gemma cups in 2-week-old plants. Scale bars represent 100 µm. **(P–Y)** Magnified views of toluidine-blue-stained sections of gemma cups in 2-week-old plants. Arrowheads and arrows indicate gemma initials and mucilage papillae, respectively. Scale bars represent 10 µm.

### Molecular characterization of *kar* mutants

The segregation ratio of the mutant gemma phenotype in an F_1_ population generated from a cross between the *kar-1* mutant, which is a female, and the wild-type (WT) male accession Takaragaike-1 (Tak-1) was 102:108 (*kar*:WT). This fit the expected 1:1 ratio as indicated by the chi-squared test (p<0.01), suggesting the involvement of a single genetic locus in the *kar* phenotype. However, the *kar* phenotype in F_1_ progenies segregated independently from the hygromycin-resistant marker in the transformed T-DNA fragment, indicating that the *kar-1* mutation is independent of the T-DNA insertion. The *kar-2* mutant, which is a male line, was infertile in several attempts at crossing with the wild-type female accession Takaragaike-2 (Tak-2).

To identify the causal gene of the *kar-1* and *kar-2* mutant phenotype, we sequenced the whole genomes of these mutants by next-generation DNA sequencing, and mapped the obtained reads on the reference genome of *M. polymorpha* [9]. Compared to the wild type, the *kar-1* mutant carried a 9-bp deletion and an 18-bp insertion at the junction of the 5th exon and 5th intron of *Mapoly0171s0028* [9]. As a result, the cDNA sequence of *Mapoly0171s0028* in the *kar1* mutant showed an 11-bp deletion and a 1-bp insertion in the coding sequence, which caused a frame-shift and generated a truncated protein (Figure S2A–C). Furthermore, we identified a deletion of an approximately 20-kb genomic locus containing the entire coding sequence of *Mapoly0171s0028* in the *kar-2* mutant (Figure S2D and E).

We then performed genetic complementation tests in the *kar-1* and *kar-2* mutants by introducing the *Mapoly0171s0028* cDNA fragment under the control of its own promoter (*proKAR:KAR*). The resultant transgenic lines in the *kar-1* and *kar-2* mutant backgrounds had restored gemma formation (Figure 1I, J, N, O, S, and T). For further confirmation, we disrupted *Mapoly0171s0028* in the wild type using homologous recombination-mediated gene targeting [11] and isolated two independent knockouts of *Mapoly0171s0028* (Figure 2A and B). The two knockout lines, *kar^KO^* #1 and #2, showed a complete loss of gemma formation similar to the *kar* mutants. The impaired gemma formation was recovered by the introduction of citrine-fused wild-type cDNA of *Mapoly0171s0028* (Figure 2C–E). These results indicated that the *kar* phenotype was caused by a loss of function of *Mapoly0171s0028*. This gene was designated as *KARAPPO* (*KAR*) after a Japanese word meaning “empty”, representing the characteristic phenotype of the mutants with empty gemma cups.

**Figure 2.**
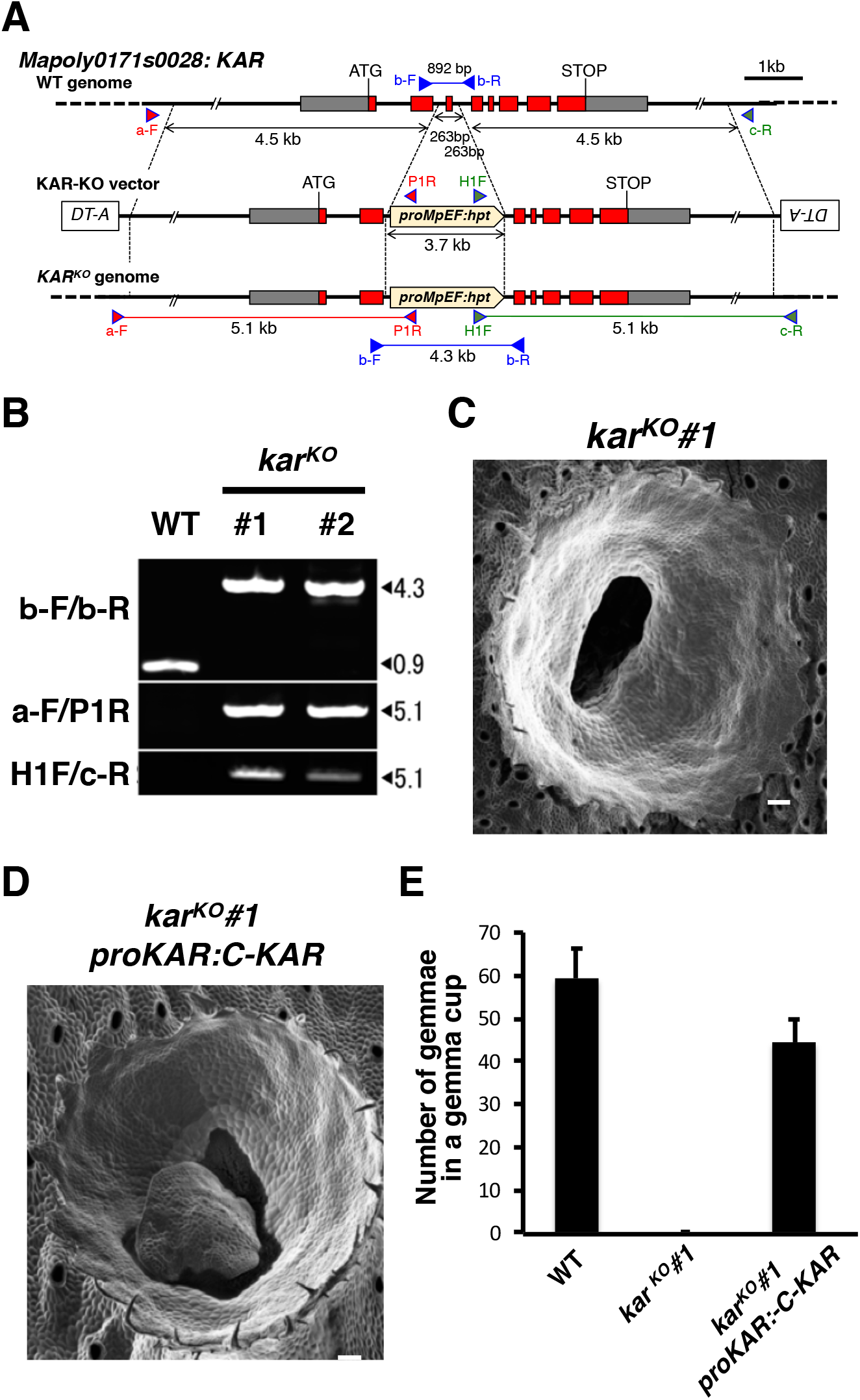
**Generation of knockout mutants of *KAR*** Schematic representation of the structure of the wild-type *KAR* locus (top), the construct designed for gene targeting (middle), and the *KAR* locus disrupted in the gene-targeted lines (bottom). Each primer pair used for genotyping is indicated by arrowheads and marked with (F) for forward or (R) for reverse. Open boxes indicate exons. Genomic PCR analysis of the *KAR^KO^* lines using the primers indicated in (A). **(C, D)** Scanning electron microscopy of gemma cups in three genotypes are presented: *kar^KO^#1* **(C)**, and a representative *kar^KO^* line transformed with *proKAR:C-KAR*, which contains a citrine-fused *KAR* coding sequence under the endogenous *KAR* promoter. **(D)**. Scale bars represent 100 µm. **(E)** Number of gemmae formed in a gemma cup in 3-week-old thalli grown from apical fragments in the wild type, a *kar^KO^* line, and a representative *kar^KO^* complemented line (Values are means ± SD, n = 5).

### *KAR* encodes a potential activator of Rop GTPase signaling

The deduced amino acid sequence of KAR encodes a highly conserved plant-specific Rop nucleotide exchanger (PRONE) catalytic domain, which is characteristic of the guanine nucleotide exchange factor (GEF) of the Rop GTPase [19], while the N- and C-terminal regions outside of the PRONE domain were highly variable (Figure S3). In angiosperms, Rop signaling mediated by RopGEF is involved in various developmental processes and environmental responses [20–25]. *KAR* is the sole PRONE-type RopGEF gene in the *M. polymorpha* genome. In addition, the *M. polymorpha* genome also contains only a single copy of *Rop*, *Mapoly0051s0092*, designated as Mp*Rop*, which showed a high overall similarity to *Rop* in various plant lineages (Figure S4A and C).

To determine whether *KAR* encodes a functional GEF that acts on MpRop, we examined the interaction of KAR and MpRop by a yeast two-hybrid assay. Yeast cells co-transformed with a combination of either AD::KAR and BD::MpRop or AD::MpRop and BD::KAR grew on selective –W/L/H and –W/L/H/A medium (Figure 3A), indicating that KAR and MpRop physically interact. The interaction between KAR and MpRop was further confirmed by an *in vitro* pull-down assay. The predicted protein coding sequence of KAR was fused to the C-terminus of the 6x Histidine-tag. Purified 6xHis-KAR fusion proteins were pulled down with guanosine triphosphate (GTP)-bound, guanosine diphosphate (GDP)-bound, or nucleotide-free forms of the glutathione S-transferase (GST)-MpRop fusion protein and were detected using anti-His antibody. KAR fusion proteins exhibited similar interactions with different forms of MpRop *in vitro* (Figure 3B).

**Figure 3.**
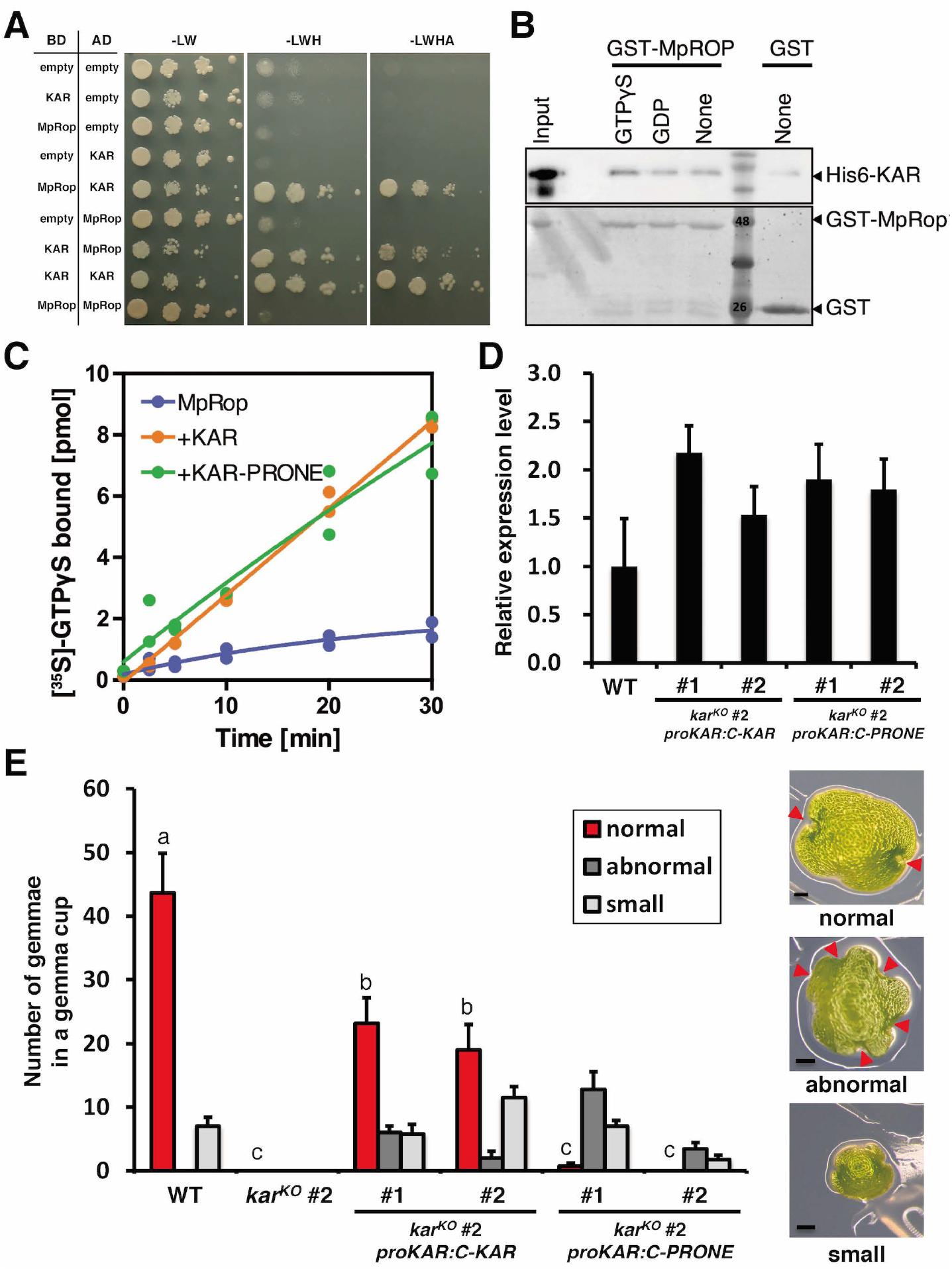
**KAR has GEF activity towards MpRop. (A)** Yeast two-hybrid experiments. Clones in the pGBKT7 vector containing the Gal4-binding domain (BD) are noted in the left column and clones in the pGADT7 vector containing the Gal4 activation domain (AD) are noted in the right column. Growth with serial dilutions on the –L, –W dropout media indicates that both pGBKT7 and pGADT7 vectors were present. Growth with serial dilutions on the –L, –W, –H dropout media and –L, –W, –H, –A dropout media indicates a physical interaction between the BD and AD fusion proteins. **(B)** Physical interaction of KAR with MpRop. His6-KAR was pulled down with GST or GST-MpRop1 using glutathione agarose. The pull-down samples were separated on a SDS-PAGE gel, then visualized by western blot with an anti-6xHis antibody. **(C)** GEF activity of the full-length KAR (KAR) or the PRONE domain of KAR (KAR-PRONE) toward MpRop. [^35^S]-GTPγS binding to 1 µM GST-MpRop1 was analyzed over time at 4°C. Graphs show data from two experiments. Fitting curves were estimated by the one-phase association model in GraphPad Prism software. **(D)** RT-qPCR analysis of *KAR* or *KAR-PRONE* expression in 3-week-old wild type and the respective complementation lines shown in Figure 3E. Mp*APT* was used as a control gene. Data are displayed as means ± SD (n = 3). **(E)** Genetic complementation with the full-length *KAR* and the truncated *KAR* coding sequence containing only the PRONE domain. The histogram shows distribution of the different classes of gemmae in a gemma cup in 3-week-old thalli grown from apical fragments, in WT, a *kar^KO^* mutant, and two representative *kar* complementation lines with the full-length KAR (*proKAR:C-KAR*) and with the PRONE domain (*proKAR:C-PRONE*) (Values are means ± s.d., n=4~5). Tukey’s test was performed for the number of normal gemmae, and letters above the bars indicate significant differences at *p* < 0.05. Right panels show pictures of normal gemma of over 500 µm diameter (normal), abnormal gemma with more than two notches, and small gemma of less than 500 µm diameter.

Next, we examined the GEF activity of KAR toward MpRop (Figure 3C). In *Arabidopsis thaliana*, there are 14 RopGEFs with a high degree of sequence similarity to the residues that are involved in catalyzing GDP/GTP exchange [26]. In Arabidopsis, the PRONE domain is sufficient for catalysis of nucleotide exchange on ROP [19] [26], while the variable C-terminal domain of some RopGEFs autoinhibits GEF activity [26]. To test the GEF activity and the potential regulatory role in the variable regions of KAR, we purified the full-length KAR protein and a truncated version of KAR containing just the PRONE domain (KAR-PRONE), and characterized their GEF activity toward MpRop using radio-labelled [^35^S]-GTPγS. We detected significant GEF activities toward MpRop with full-length KAR and KAR-PRONE, and the GEF activity of full-length KAR was comparable to that of KAR-PRONE (Figure 3C).

We tested the functionality of the *KAR-PRONE* coding sequence without the N- and C-terminal region in gemma formation. Two lines with comparable expression levels of the full-length *KAR* or the *KAR-PRONE* coding sequence were selected for further analysis (Figure 3D). Introduction of citrine-fused full-length *KAR* cDNA restored to some extent the formation of gemma with normal morphology in the *kar^KO^* #2 background. By contrast, the citrine-fused KAR-PRONE did not lead to the full recovery of gemma formation; the most of the gemmae were abnormal in size and morphology with an irregular periphery (Figure 3E). These results demonstrated that the N-terminal and C-terminal variable regions of KAR play a significant role in proper gemma formation *in viv*o, although they have no obvious effect on GEF activity in vitro (Figure 3C).

### Ubiquitous expression of *KAR* and Mp*Rop* in vegetative tissues

To evaluate the expression pattern of *KAR* and Mp*Rop*, we generated transgenic *M. polymorpha* lines expressing the -glucuronidase (*GUS*) reporter gene under the control of the *KAR* promoter (*proKAR:GUS*) and the Mp*Rop* promoter (*pro*Mp*Rop:GUS*). In *proKAR:GUS* and *pro*Mp*Rop:GUS* lines, GUS staining was observed in the broader region of the entire thallus, including the basal floor of the gemma cups containing developing gemmae (Figure 4A and B). We further evaluated the expression pattern of *KAR* and Mp*Rop* using reverse-transcription quantitative PCR (RT-qPCR). Transcripts of *KAR* and Mp*Rop* were detected in all stages and organs in the vegetative thallus (Figure 4C). These results suggest that *KAR* and Mp*Rop* are ubiquitously and simultaneously expressed in the initial stage of gemma development.

**Figure 4.**
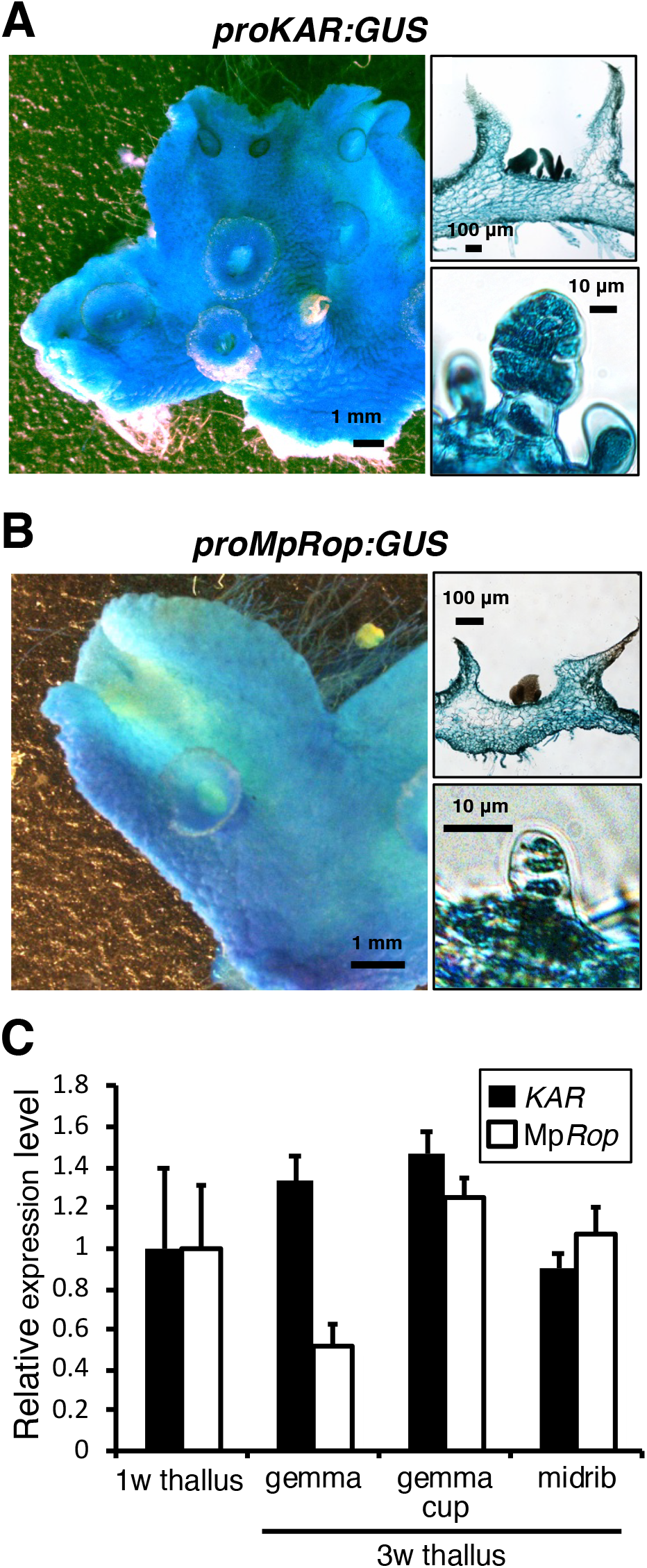
**Expression of KAR and MpRop in vegetative tissues (A, B)** Histological GUS activity staining of representative proKAR:GUS **(A)** and proRop:GUS **(B)** transgenic lines. **(C)** RT-qPCR analysis of *KAR* and Mp*Rop* in 1-week-old thalli yet to develop gemma cups (1w thallus), mature gemmae in gemma cups (gemma), gemma cups containing developing gemmae (gemma cups), and midribs. Total RNA was isolated from the respective tissues of Tak-1. *EF1α* was used as a control gene. Data are displayed as means ± SD (n = 3).

Knockout plants of Mp*Rop* were generated by homologous recombination-mediated gene targeting; however, two independent Mp*Rop^KO^* lines showed severe impairment of thallus growth and wilted before the formation of the gemma cup (Figure S4B). This result suggests a much broader function for the sole Rop in *M. polymorpha*, which is evidently essential for the growth and development of the gametophyte thallus. Another type of GEF, SPIKE1 (SPK1), has been reported to regulate cytoskeletal rearrangement and cell-shape change in response to growth signals in angiosperms [27, 28]. SPK1 has a conserved DOCK homology region 2 (DHR2) domain, which is distantly related to CZH [CDM (Ced-5, Dock180, Myoblastcity)-Zizimin homology] RhoGEFs in animals and fungi [29]. *M. polymorpha* contains a single *SPK1* homologue, Mp*SPK1* [9] (Figure S4C), which may have a critical function in controlling Rop signaling in thallus growth.

The contrast between the specific developmental impairment in *kar* mutant lines (Figure 1 and 2), and the ubiquitous promoter activity of *KAR* and Mp*Rop* (Figure 4), suggests an upstream regulatory mechanism for KAR activity, which enables cell-type specific activation of Rop in the gemma initial. Recent studies have shown that several plasma membrane-localized receptor-like protein kinases (RLKs) function as upstream regulators of Rop signaling through interaction with the C-terminal variable region of RopGEF (*i.e.* the pollen receptor kinases in tomato (*Solanum lycopersicum*) and FERONIA in *Arabidopsis thaliana*) [20, 21, 30]. Similar to their role in polarized cell growth mediated by Rop signaling in angiosperms, RLK(s) might be involved in the specific function of KAR in the initial stage of gemma development in *M. polymorpha*. Further functional studies on the N-terminal and C-terminal variable regions of KAR will be needed to understand the regulatory mechanism of Rop signaling in *M. polymorpha*.

### Role of KAR in the initial stage of gemma development

In this study, we demonstrated that *KAR*, which encodes a RopGEF, is an essential factor required for the initial stage of gemma development in *M. polymorpha*. Histological studies have suggested the occurrence of cell protrusion and subsequent asymmetric cell divisions in the initial stage of gemma development in the epidermal floor of the gemma cup (Figure S1; [2]). A recent study in *M. polymorpha* demonstrated that a ROOT-HAIR DEFECTIVE SIX-LIKE (RSL) class I basic helix-loop-helix (bHLH) transcription factor, MpRSL1, controls the morphogenesis of structures derived from individual epidermal cells (*i.e.* rhizoids, slime papillae, mucilage papillae, and gemmae) [3]. Similar to the loss-of-function mutants of Mp*RSL1*, in the *kar* mutants, we did not observe the one- or two-cell stage of gemma development. On the other hand, the other epidermis-derived structures, mucilage papillae and rhizoids, were generated normally in the *kar* mutants, but are absent in Mp*rsl1* mutants [3]. Mucilage papillae and rhizoids do not undergo any further cell division after polarized cell growth, whereas the development of gemmae involves subsequent asymmetrical cell division and further cell divisions. These observations suggest that KAR promotes the asymmetrical cell division(s) of the gemma precursor cell after the specification of an epidermal cell controlled by MpRSL1.

KAR contains a highly conserved PRONE domain (Figure S3), which has been implicated in the activity of the GEF of Rop GTPase [19, 26]. In this study, we demonstrated the GEF activity of KAR on MpRop *in vitro* (Figure 3). The *Arabidopsis thaliana* genome contains 14 RopGEFs, and Rop signaling mediated by RopGEF is involved in the control of polar cell growth of pollen tubes and root hairs [20–23]. This polar growth involves the coordination of cytoskeleton organization and vesicular trafficking [31, 32]. In yeast and animals, the closest homologues of Rop, the Cdc42 Rho GTPases, regulate polarity and play a key role in the control of asymmetrical cell division [33, 34]. Recently in monocots, Rop was shown to be involved in the asymmetrical division of the stomata mother cell by controlling cytoskeletal scaffolds and nuclear positioning [35, 36]. KAR-mediated Rop signaling could function in the formation of cell polarity and/or subsequent asymmetrical cell divisions during the initiation of gemma development.

Vegetative reproduction can be considered as a type of naturally occurring somatic embryogenesis, in which a meristem is regenerated from differentiated cells. In the development of somatic embryos from single cells isolated from tissue cultures of carrot (*Daucus carota* subsp. *sativus*), the first cell division occurs asymmetrically, and one of the daughter cells gives rise to a three-dimensional cell mass from which one or more embryos develop [4–6]. Asymmetrical cell division to produce daughter cells of a different cell fate must be a common key process in the initial stage of organ/plantlet regeneration from differentiated cells. RopGEF-mediated Rop signaling seems to have been acquired in the common ancestor of land plants after the emergence of charophycean algae. PRONE-type RopGEFs have highly diverged in the course of land plant evolution, while the basal land plant *M. polymorpha* has a limited gene repertoire for Rop signaling (Figure S3 and S4). The Rop-driven asymmetric cell division of differentiated cells to regenerate clonal progenies could be a key innovation for sessile land plants to dominate the terrestrial ecosystem, and this mechanism may have been co-opted to regulate numerous physiological and developmental processes, probably also including vegetative reproduction or organ regeneration, in the course of land plant evolution.

## Contact for Reagent and Resource Sharing

Further information and requests for resources and reagents should be directed to and will be fulfilled by the Lead Contact, Kimitsune Ishizaki. Please note that the transfer of transgenic plants will be governed by an MTA, and will be dependent on appropriate import permits being acquired by the receiver.

## Supplemental Information

Figures S1–S4, and Table S1.

## Author Contributions

T.H. and K.I. designed the research, and T.H. performed most of the experiments. K.I. isolated the *kar-1* mutant and H.K. performed linkage analyses. M.K. and K.T.Y. isolated *kar-2* mutants. T.H., Y.Y., and H.T. performed the histology. K.L.Q. and D.U. performed the *in vitro* pull-down and GEF assays. K.I., K.Y., S.Sh., S.Sa., and T.K. performed the whole-genome sequencing. M.S., M.W., and K.T. performed TEM analyses. T.H., K.I., and T.K. performed the gene-targeting experiments. T.H., K.I., Y.Y., D.U., H.F., and T.M. analyzed the data. T.H., Y.Y., and K.I. wrote the article.

## Acknowledgments

The authors thank Shohei Yamaoka and Ryuichi Nishihama for critically reading the manuscript; Tatsuaki Goh, Kohichi Toyokura, and Miwa Ohnishi for discussions; Sakiko Ishida, Yoriko Matsuda, and Chiho Hirata for technical assistance. Whole-genome sequencing was supported by NIBB Collaborative Research Programs (15-823 to K.I.). This study was supported by MEXT KAKENHI Grants-in-Aid for Scientific Research on Innovative Area (25113009 to T.K., and 25119711, 25114510, and 17H06472 to K.I.), JSPS KAKENHI Grants-in-Aid for Scientific Research (B) (15H04391 to K.I.), and by the Asahi Glass Foundation and SUNTORY Foundation for Life Sciences (to K.I.).

## Star Methods

## Key Resources Table

(Attached)

## Contact for Reagent and Resource Sharing

Further information and requests for resources and reagents should be directed to and will be fulfilled by the Lead Contact, Kimitsune Ishizaki. Please note that the transfer of transgenic lants will governed by an MTA, and will be dependent on appropriate import permits being acquired by the receiver.

## Experimental Modeland Subject Details

### Plant Materials and Growth Conditions

Female and male accessions of *M. polymorpha*, Takaragaike-2 (Tak-2) and Takaragaike-1 [10], respectively (Ishizaki et al. 2008), were used as the wild type. F_1_ spores generated by crossing Tak-2 and Tak-1 plants were used for transformation to generate the *kar-1* mutant and the *kar* knockout lines. Thalli were grown on 1% (w/v) agar medium containing half-strength Gamborg’s B5 salts [37] under 50–60 µmol m^−2^ s^−1^ continuous white light with a cold cathode fluorescent lamp (CCFL; OPT-40C-N-L; Optrom, Japan) or white light-emitting diodes (white LED; VGL-1200W; SYNERGYTEC, Japan) at 22°C. For crossing, over 2-week-old thalli were transferred to continuous light conditions with 50–60 µmol m^−2^ s^−1^ white LED and 20–30 µmol m^−2^ s^−1^ far red light-emitting diodes (VBL-TFL600-IR730, Valore, Japan).

## Method Details

### Phenotype Analysis and Histology

Two-week-old thalli developed from tips of thalli were dissected into small pieces and transferred to fixative solution with 2% glutaraldehyde in 0.05 M phosphate buffer (pH 7.0) and evacuated with a water aspirator until the specimens sank, then fixed for 2 days at room temperature. The samples were dehydrated in a graded ethanol series and embedded in Technovit 7100 plastic resin. Semi-thin sections (5-μm thickness) were obtained with a microtome (HM 335E*, Leica* Microsystems) for light microscopy and stained with toluidine blue O. Sections were observed with an upright microscope (Axio Scope. A1, Carl Zeiss Microscopy).

Cultured thalli of *M. polymorpha* were observed using a digital microscope (VHX-5000, KEYENCE). For scanning electron microscopy, thalli were frozen in liquid nitrogen and observed with a scanning electron microscope (VHX-D500, KEYENCE).

### Electron microscopy

Samples were fixed with 4% paraformaldehyde and 2% glutaraldehyde in 50 mM sodium cacodylate buffer (pH 7.4) for 2 h at room temperature and overnight at 4°C, then post-fixed with 1% osmium tetroxide in 50 mM cacodylate buffer for 3 h at room temperature. After dehydration in a graded methanol series (25, 50, 75, 90, and 100%), the samples were embedded in Epon812 resin (TAAB). Ultrathin sections (100 nm) or semi-thin sections (1 μm) were cut by a diamond knife on an ultramicrotome (Leica EM UC7, Leica Microsystems, Germany) and placed on a glass slide. The sections were stained with 0.4% uranyl acetate followed by lead citrate solution and coated with osmium under an osmium coater (HPC-1SW, Vacuum Device, Japan). The coated sections were observed with a field-emission scanning electron microscope SU8220 (Hitachi High technology, Japan) with an yttrium aluminum garnet backscattered electron detector at an accelerating voltage of 5 kV.

### Genome sequencing of the *kar-1* and *kar-2* mutants

Genomic DNA was extracted from *kar-1, kar-2*, and Tak-2 plants as follows: the tissue was powdered in liquid nitrogen and incubated in 10 mL hexadecyltrimethylammonium bromide (CTAB) buffer (1.5% CTAB, 75 mM Tris-HCl [pH 8.0], 15 mM EDTA, and 1 M NaCl) for 20 min at 56°C. This suspension was mixed with an equal volume of chloroform:isoamyl alcohol (24:1, w/v), incubated for 20 min at room temperature, and centrifuged at 4,000 × g for 20 min. The aqueous phase was used to repeat the chloroform:isoamyl alcohol extraction, and then mixed gently with 1.5 volumes of CTAB precipitation buffer (1% CTAB, 50 mM Tris-HCl [pH 8.0], and 10 mM EDTA). After centrifugation at 10,000 × g for 30 min at 20°C, the precipitate was dissolved in 1 M sodium chloride containing 10 mg/mL RNaseA and incubated for 30 min at 37°C. The genomic DNA was precipitated with ethanol, dissolved in TE buffer (10 mM Tris-HCl [pH 8.0] and 1 mM EDTA), and further purified using a Genomic-tip 100 column (QIAGEN, Germany). The DNA was then sheared on a Covaris sonicator (Covaris, USA), size-selected with Pippin Prep (Sage Science, USA), and used to create the libraries using the TruSeq DNA Sample Preparation Kit (Illumina) with an insert size of ~350 bp. The libraries were sequenced using Illumina HiSeq 2000 with a 2 × 101-nt paired-end sequencing protocol. The sequence reads were mapped to the *M. polymorpha* genome sequence [9] and the plasmid sequence was used for the biolistic transformation [18] by Bowtie2 v.2.2.9 [38] with default parameters, and visualized and assessed using Integrative Genomics Viewer v.2.3.23 [39].

### Characterization of mutations in *KAR/Mapoly0171s0028*

Small pieces (3 × 3 mm) of thalli were taken from individual plants and crushed with a micro-pestle in 100 µl buffer containing 100 mM Tris-HCl, 1 M KCl, and 10 mM EDTA (pH 9.5). Sterilized water (400 µl) was added to each tube and a 1 µl aliquot of the extract was used as a template for PCR using KOD FX Neo DNA polymerase (Toyobo). To identify the mutation in the *kar-1* mutant, the *Mapoly0171s0028* locus was amplified by genomic PCR using the primer set kar-1_gF/kar-1_gR and sequenced. The cDNA of *kar* in the *kar-1* mutant was amplified by RT-PCR using the primer set KAR-cds-F/KAR-cds-sR and sequenced. To confirm the absence of *Mapoly0171s0028* in the *kar-2* mutant, genomic PCR was performed with a KOD FX Neo DNA polymerase using primers KAR-gF/KAR-gR. Primer pairs are shown in Figure S2D and Table S1.

### Complementation tests

For complementation of the *kar* mutants, the coding sequence of full-length *KAR* and the truncated coding sequence of *KAR* (*KAR-PRONE*), containing just the PRONE domain (residues 132–503) were amplified by RT-PCR using KOD plus neo (TOYOBO) with the primer set KAR-cds-F/KAR-cds-sR and PRONE-L/PRONE-R, respectively. The *KAR* and *KAR-PRONE* coding sequence fragments were cloned into the pENTR/D-TOPO cloning vector (Life Technology) to produce pENTR-KAR and pENTR-PRONE, respectively. The *KAR* promoter region, including about 5 kb upstream of the initiation codon, was amplified from Tak-1 genomic DNA by PCR using KOD-Plus-Neo (TOYOBO) with the primer set KARpro_GW_F/KARpro_GW_302_R. The PCR-amplified product was cloned into the *Xba*I and *HindⅢ* sites of pMpGWB302 to replace the CaMV35S promoter [40] with the In-Fusion HD cloning kit (Clontech, Mountain View, CA). The entry vector containing the *KAR* coding sequence was introduced into the binary vector by Gateway LR clonase II Enzyme mix (Thermo Fisher Scientific, USA) to generate the *proKAR:KAR* construct. The *proKAR:KAR* vector was introduced into regenerating thalli of *kar-1* and *kar-2* mutants via *Agrobacterium tumefaciens* GV2260 [13].

Similarly, the *KAR* promoter region, including about 5 kb upstream of the initiation codon, was amplified from Tak-1 genomic DNA by PCR using KOD-Plus-Neo (TOYOBO) with the primer set KARpro_GW_F/KARpro_GW_305_R. The PCR-amplified product was cloned into the *Xba*I and *HindⅢ* sites of pMpGWB305, which contains citrine gene in front of the gateway cassette, to replace the CaMV35S promoter [40] with the In-Fusion HD cloning kit (Clontech, Mountain View, CA). The coding sequence fragments of *KAR* and *KAR-PRONE* in the entry vectors pENTR-KAR and pENTR-PRONE were introduced into the binary vectors by Gateway LR clonase II Enzyme mix (Thermo Fisher Scientific, USA) to generate the *proKAR:C-KAR*, and *proKAR:C-PRONE* constructs, respectively. The binary vector was transformed into the *kar^KO^* line. Transformants were selected with 0.5 µM chlorsulfuron and 100 µg/ml cefotaxime.

### Generation of *KARKO* and Mp*RopKO* plants

To generate the *KAR*-targeting vector, 5′- and 3′-homologous arms (approximately 4.5-kb in length) were amplified from Tak-1 genomic DNA by PCR using KOD FX Neo (TOYOBO) with the primer pairs shown in Table S1. The PCR-amplified 5′- and 3′-homologous arms were cloned into the *Pac*I and *Asc*I sites, respectively, of pJHY-TMp1 [11] with the In-Fusion HD cloning kit (Clontech, Mountain View, CA). The *KAR*-targeting vector was introduced into F_1_ sporelings derived from sexual crosses between Tak-1 and Tak-2 via *Agrobacterium tumefaciens* GV2260 [10]. The transformed plants carrying the targeted insertions were selected by genomic PCR with a KOD FX Neo DNA polymerase and primer pairs shown in Figure 2 and Table S1.

To generate the Mp*Rop*-targeting vector, 5′- and 3′-homologous arms (approximately 4.5-kb in length) were amplified from Tak-1 genomic DNA shown in Table S1. The PCR-amplified 5′- and 3 ′-homologous arms were cloned into the *Pac*I and *Asc*I sites, respectively, of pJHY-TMp1 [11] with the In-Fusion HD cloning kit (Clontech, Mountain View, CA). The Mp*Rop*-targeting vector was transformed into F_1_ sporelings derived from sexual crosses between Tak-1 and Tak-2 as described above. The transformed plants carrying the targeted insertions were selected by genomic PCR with a KOD FX Neo DNA polymerase and primer pairs shown in Figure S4B and Table S1.

### Promoter reporter analyses

The *KAR* genomic sequence, including approximately 5 kb upstream of the initiation codon, was amplified from Tak-1 genomic DNA by PCR using KOD-Plus-Neo (TOYOBO) with the primer set KARpro_F/KARpro_R and was cloned into pENTR/D-TOPO (Thermo Fisher Scientific). Similarly, the Mp*Rop* genomic region, including about 3 kb upstream of the initiation codon, was amplified from Tak-1 genomic DNA by PCR with the primer set MpRoppro_F/MpRoppro_R and was inserted into pENTR/D-TOPO (Thermo Fisher). These entry vectors were introduced into the Gateway binary vector pMpGWB104 [40] using Gateway LR clonase II Enzyme mix (Thermo Fisher Scientific, USA) to generate *proKAR:GUS* and *proMpRop:GUS* binary constructs, respectively. The *proKAR:GUS* and *proMpRop:GUS* vectors were introduced into regenerating thalli of Tak-1 via *Agrobacterium tumefaciens* GV2260 [13]. Transformants were selected with 0.5 µM chlorsulfuron and 100 µg/ml cefotaxime. For histological GUS activity assays, transformants were incubated in GUS staining solution (0.5 mM potassium ferrocyanide, 0.5 mM potassium ferricyanide, and 1 mM X-Gluc) at 37°C and later cleared with 70% ethanol (Jefferson et al., 1987).

### Yeast Two-Hybrid (Y2H) assay

To construct AD::KAR and AD::MpRop vectors, the *KAR* and Mp*Rop* coding sequences were amplified by PCR using KOD plus neo (TOYOBO) with primer pairs KAR_WT_Y2H_pGADT7_F and KAR_WT_Y2H_pGADT7_R, and Rop_WT_Y2H_pGADT7_F and Rop_WT_Y2H_pGADT7_R, respectively, and subcloned into the *Not*I site of pGADT7-AD in Matchmaker Gold Yeast Two-Hybrid System (Takara Bio, Japan) with the In-Fusion HD cloning kit (Takara Bio). To construct BD::KAR and BD::MpRop vectors, the *KAR* and Mp*Rop* coding sequences were amplified by PCR using KOD plus neo (TOYOBO) with primer pairs KAR_WT_Y2H_pGBKT7_F and KAR_WT_Y2H_pGBKT7_R, and Rop_WT_Y2H_pGBKT7_F and Rop_WT_Y2H_pGBKT7_R, respectively, and subcloned into the NotI site of pGBKT7 in Matchmaker Gold Yeast Two-Hybrid System (Takara Bio) with the In-Fusion HD cloning kit (Takara Bio). Indicated combinations of plasmids were co-transformed into yeast strain Y2H Gold (Takara Bio) following the protocol for high-efficiency transformation of yeast with lithium acetate, single-stranded carrier DNA, and polyethylene glycol. Following transformation, colonies were selected for the presence of the plasmids, inoculated in liquid synthetic drop-out (SD) media (lacking the amino acids leucine and tryptophan, with the exception of untransformed strain Y2H Gold, which was grown in YPD), grown to saturation, and plated onto SD media plates lacking the indicated amino acids. SD media plates lacked the amino acids leucine, tryptophan, and histidine (SD –Leu/–Trp/–His). Serial 1:5 dilutions were made in water and 3 μl of each dilution was used to yield one spot. Plates were incubated at 30°C for two (SD –Leu/–Trp) or three (SD –Leu/–Trp/–His) days before taking pictures.

## Protein purification

The cDNA of MpRop was amplified by RT-PCR using KOD plus neo (TOYOBO) with primer pairs MpRop-cds-F and MpRop-cds-sR and cloned into pENTR/D-TOPO (Thermo Fisher Scientific). The Mp*Rop* coding sequence in the resultant ENTRY clone and pENTR-KAR and pENTR-PRONE generated above were transferred using Gateway LR clonase II Enzyme mix (Thermo Fisher Scientific, USA) into the bacterial expression vectors pDEST15 or pDEST17, which express GST- or 6xhistidine (6xHis)-tagged protein, respectively. 6xHis-KAR and 6xHis-KAR-PRONE were expressed in the *Escherichia coli* strain Arctic Express RP with 0.25 mM IPTG at 12°C. The cells were harvested by centrifugation and lysed in extraction buffer (20 mM Tris-HCl (pH 7.4), 200 mM NaCl, 2 mM MgCl_2_, 5 mM 2-mercaptoethanol, 1 mM phenylmethane sulfonyl fluoride or phenylmethylsulfonyl fluoride (PMSF), 2 µg/ml leupeptin, 250 µg/ml lysozyme, and 0.2% C_12_E_10_) with sonication. The bacterial lysate was centrifuged at 100,000 x g for 1 hr. His-tagged protein was captured from the supernatant using nickel-NTA agarose, washed with wash buffer (20 mM Tris-HCl (pH 7.4), 500 mM NaCl, 5 mM 2-mercaptoethanol, 1 mM PMSF, 2 µg/ml leupeptin, and 20 mM imidazole) and eluted with elution buffer (20 mM Tris-HCl (pH 7.4), 150 mM NaCl, 5 mM 2-mercaptoethanol, 1 mM PMSF, and 20% glycerol) with 250 mM imidazole. The eluted proteins were dialyzed against elution buffer and frozen at –80°C.

Bacterial cells expressing GST or GST-MpRop were lysed in extraction buffer (50 mM Tris-HCl (pH 7.4), 150 mM NaCl, 2 mM MgCl_2_, 1 mM EDTA, 5 mM 2-mercaptoethanol, 1 mM PMSF, 2 µg/ml leupeptin, 100 μM GDP, 250 µg/ml lysozyme, and 0.2% C_12_E_10_) with sonication. The bacterial lysate was centrifuged at 100,000 x g for 1 hr. GST-tagged proteins were captured from the supernatant using glutathione-agarose, washed with wash buffer I (50 mM Tris-HCl (pH 7.4), 100 mM NaCl, 1 mM EDTA, 5 mM 2-mercaptoethanol, 40 μM GDP, 2 mM MgCl_2_, 1 mM PMSF, and 2 mµ/ml leupeptin), wash Buffer II (50 mM Tris-HCl (pH 7.4), 500 mM NaCl, 40 μM GDP, 2 mM MgCl_2_, 1 mM EDTA, 5 mM 2-mercaptoethanol, 1 mM PMSF, and 2 µg/ml leupeptin), then wash buffer I again. GST-tagged proteins were eluted with elution buffer (100 mM Tris-HCl (pH 8.8), 200 mM NaCl, 40 μM GDP, 2 mM MgCl_2_,10 mM 2-mercaptoethanol, 1mM EDTA, 1 mM PMSF, and 20% glycerol) with 20 mM glutathione. The eluted proteins were dialyzed against elution buffer without GDP and frozen at –80°C.

### *In vitro* pull-down assay

One μg of 6xHis-KAR was incubated with 10 μg of GST or GST-MpROP in Nucleotide Binding Buffer (20 mM HEPES-NaOH (pH 7.4), 5 mM MgCl_2_, 1 mM dithiothreitol, 0.1% Triton X-100, and 1 mM EDTA) preloaded with 10 μM GDP, GTPγS, or no nucleotide for 4 hours at 4°C. The samples were centrifuged with a table-top centrifuge at full speed for 1 min, and the supernatant was incubated with Glutathione-agarose resin for 30 min at 4°C. The resins were then washed for three times with Nucleotide Binding Buffer with the respective nucleotide. The resins were then boiled with SDS loading buffer and separated by sodium dodecyl sulfate polyacrylamide gel electrophoresis (SDS-PAGE). The 6xHis-tagged proteins were detected by western blot with a mouse anti-6xHis antibody (Santa Cruz Biotech) as the primary antibody, an HRP-conjugated anti-mouse IgG antibody was used as the secondary antibody. GST and GST-MpRop were detected by Coomassie Brilliant Blue staining.

### GTPγS binding on MpRop

The GEF enzymatic activity of KAR or KAR-PRONE toward MpRop was analyzed using radio-labelled [35S]-GTPγS, as described in previous studies with slight modifications [26, 41]. For [^35^S]-GTPγS binding, 2 µM GST-MpRop in reaction buffer (50 mM Tris-HCl (pH 7.4), 1 mM EDTA, 1 mM DTT, and 5 mM MgCl_2_) was mixed with an equal volume of reaction buffer containing 5 µM [^35^S]-GTPγS to start the exchange reaction on ice. At given time points, 50 µl aliquots were removed and placed into 450 µl of ice-cold wash buffer (20 mM Tris-HCl (pH 7.4), 100 mM NaCl, and 25 mM MgCl_2_) with 0.1 mM GTP, then applied to a nitrocellulose membrane filter. The filter was washed three times with 2 to 3 ml of ice-cold wash buffer. The amount of [^35^S]-GTPγS was measured by scintillation counting.

### RT-qPCR

Total RNA was isolated from the 1-week-old thalli, and mature gemmae, gemma cups, and midribs of 3-week-old thalli of Tak-1 (Fig. 4C) and 1-week-old thalli of the *kar^KO^#2* line transformed with *proKAR:Citrine-KAR* or *proKAR:Citrine-PRONE* (Fig. 3D) using the RNeasy Plant mini kit (Qiagen). One μg of total RNA was reverse-transcribed in a 20 μl reaction mixture using ReverTra Ace qPCR RT Master Mix with gDNA remover (TOYOBO). After the reaction, the mixture was diluted with 40 μl of distilled water and 2 μl aliquots were used for quantitative PCR (qPCR) analysis. qPCR was performed with the Light Cycler 96 (Roche) using KOD SYBR qRT-PCR Mix (TOYOBO) according to the manufacturer’s protocol. The primers used in these experiments are listed in Table S1. Transcript levels of Mp*EF1a* or Mp*APT* were used as a reference for normalization [42]. RT-qPCR experiments were performed using three biological replicates and technically duplicated.

### Phylogenetic Analysis of KAR

For phylogenetic analysis of KAR and RopGEFs, peptide sequences were collected from genomic information of *M. polymorpha* in MalpolBase (http://marchantia.info/), *A. thaliana* in TAIR (http://www.arabidopsis.org), *Physcomitrella patens* [43] and *Selaginella moellendorffii* [44] in Phytozome (https://phytozome.jgi.doe.gov/pv/portal.html), and *Klebsormidium nitens* NIES-2285 in (http://www.plantmorphogenesis.bio.titech.ac.jp/~algae_genome_project/klebsormidium/kf_download.htm) [45]. A multiple alignment of amino acid sequences of KAR and its homologous RopGEFs was first constructed using the MUSCLE program [46] implemented in MEGA6.06 [47] with default parameters, which was performed using a Maximum Likelihood method by PhyML [48] with the LG+G+I substitution model. One thousand bootstrap replicates were performed in each analysis to obtain the confidence support.

---

**Figure S1:**
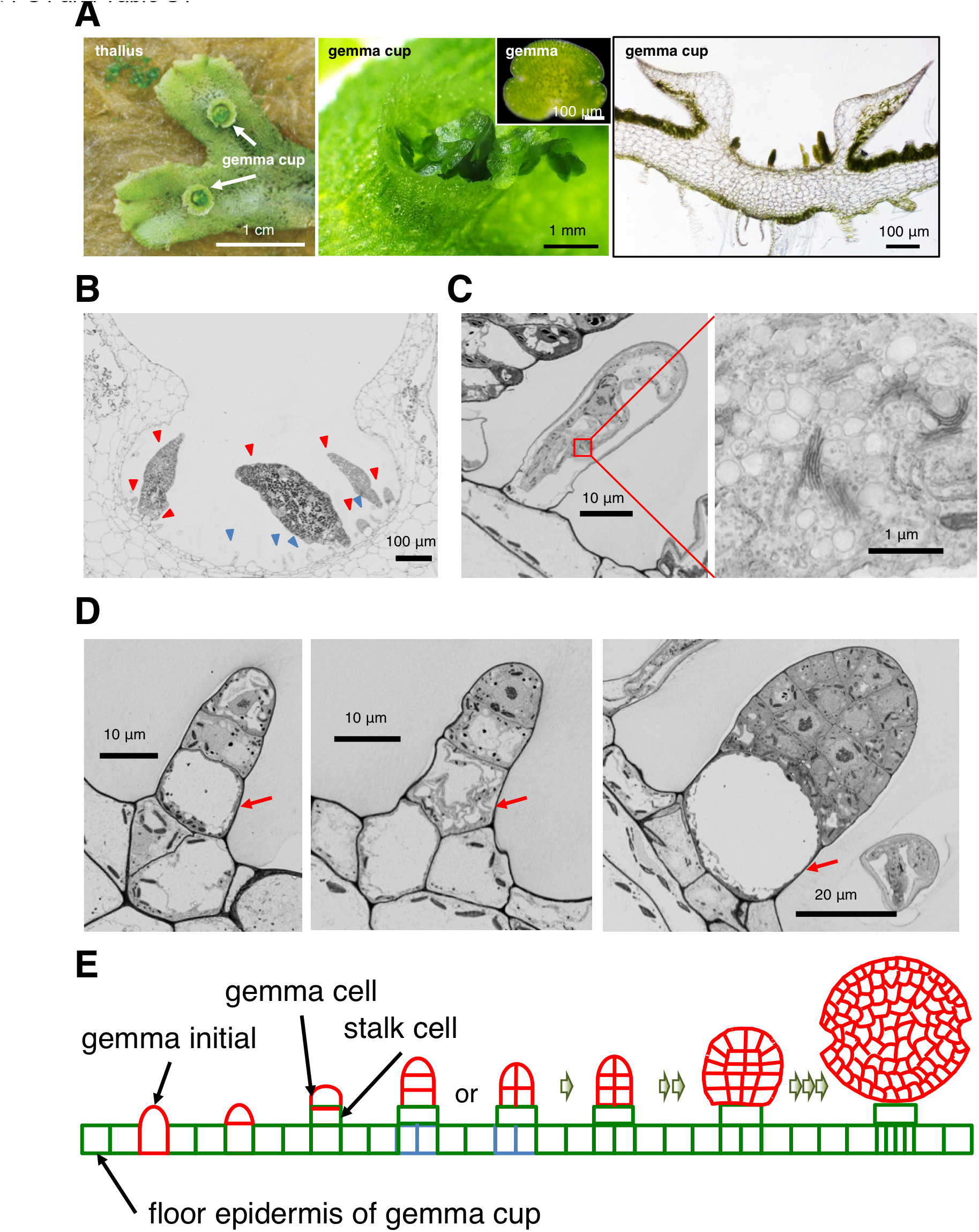
**Gemma development in *M. polymorpha*, Related to Figure 1.** (A) Optical observation of gemma and gemma cup in M. polymorpha. Gemma cups formed on the dorsal surface of gametophyte body, thallus. Top view of thallus (left), a close-up view of gemma cup (middle), and a transverse section of a gemma cup (right). (B-D) Electron microscopy in the basal floor of gemma cup. (B) Transverse section of a gemma cup. Red arrow-heads indicate developping gemmae. Blue-arrow heads indicate mucilage papillae. (C) Close up view of mucilage papillae observed at the floor epidermis of gemma cup. The right image shows an enlarged image of indicated cytosolic region in the left image. (D) Close up view of developing gemmae. Red arrows indicate basal stalk cells. (E) Schematic model of gemma development in the floor epidermis of gemma cup.

**Figure S2.**
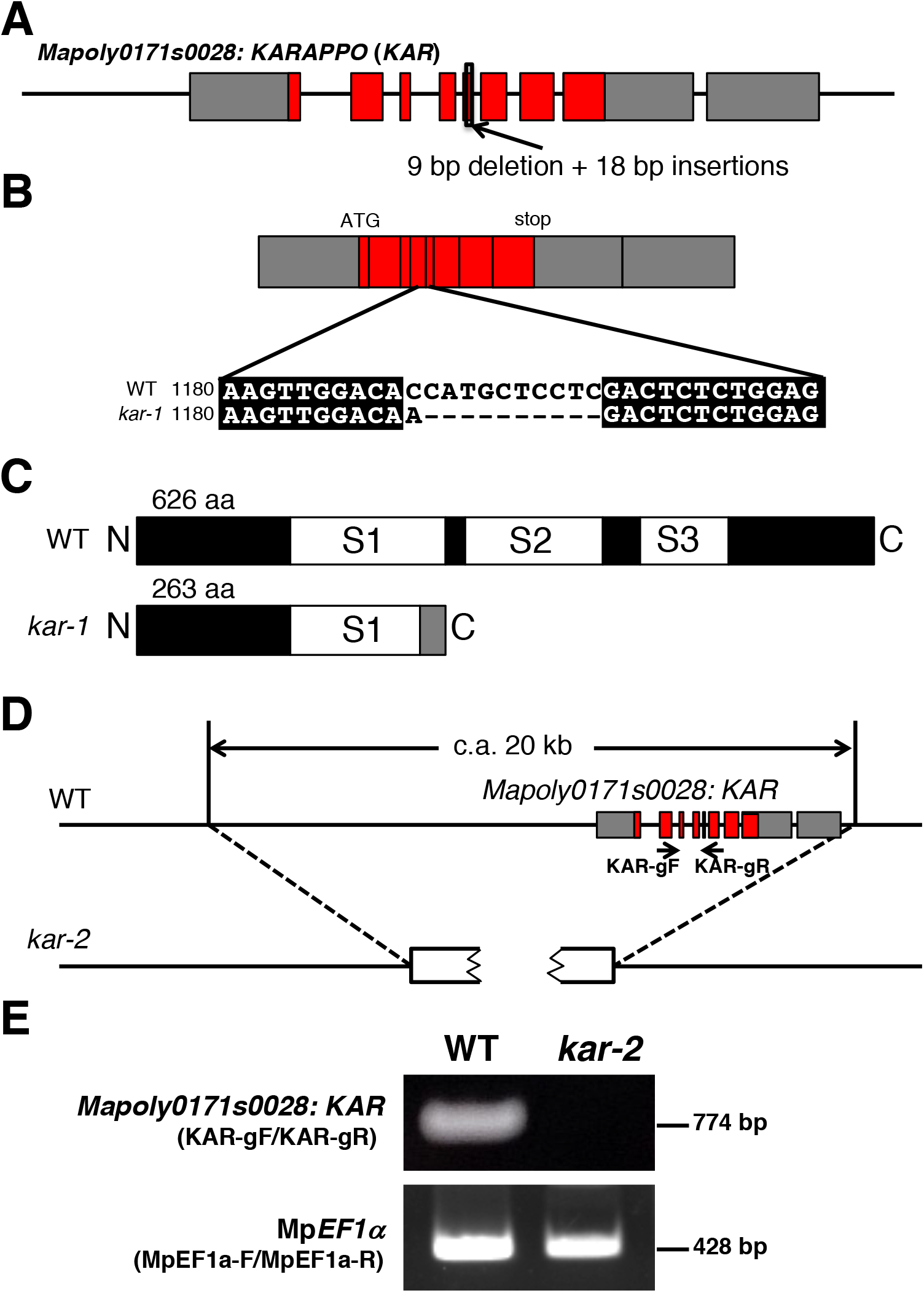
**Molecular characterization of *kar-1* and *kar-2*, Related to Figure 1 to 2.** Whole genome analysis revealed respective mutations in Mapoly0171s0028 locus in *kar-1* and *kar-2.* (A) Schematic representation of the Mapoly0178s0028 genomic locus in wild-type and *kar-1*. Gray boxes indicate exons of untranslated region. Red boxes indicate exons of protein coding region. A small deletion found in the *Mapolyo171s0028* locus of *kar-1* genome. (B) cDNA sequences of *Mapolyo171s0028* in *kar-*1*.* (C) Schematic representation of deduced gene products of *Mapolyo171s0028* in wild-type and *kar-1.* (D) Whole genome analysis revealed c.a. 20 kb deletion in *kar-2.* Broken open boxes indicate partial fragments of pMT plasmid (Takenaka et al. 2000), which was introduced by particle bombardment protocol. A series of genomic PCR suggested that the 5’ and 3’ region of Mapoly0171s0028 is not adjacent to each other in *kar-2* (data not shown), suggesting occurrence of a genomic rearrangement accompanied with the physical DNA delivery. (E) Genomic PCR of Mapoly0178s0028 in wild-type and *kar-2.* Mp*EFa; M. polymorpha* Elongation Factor alpha gene (Mp*EFa*) was used as a positive control.

**Figure S3.**
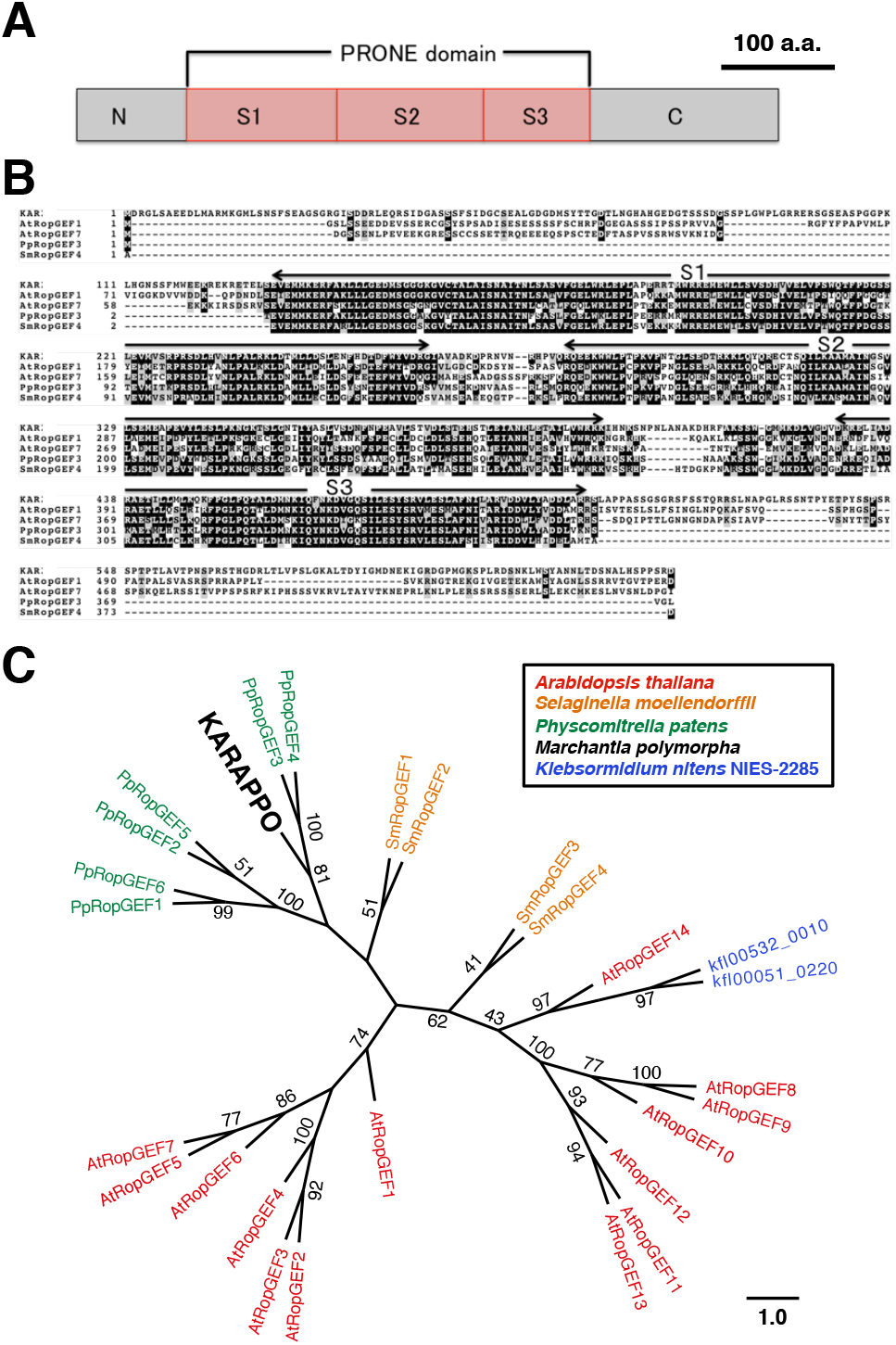
***KAR* encodes a highly conserved PRONE domain of RopGEF, Related to Figure 2 to 3.** (A) A domain structure of the *KAR* gene product. (B) Multiple alignment of the full amino acid sequences of KAR and representative RopGEFs in the moss *Physcomitrella patens*, the lycophyte *Sellaginella moellendorffii* and *Arabidopsis thaliana* RopGEFs. Lines above aligned sequences indicate highly conserved regions in a PRONE domain composed of three subdomains (S1, S2, and S3), which has been in Arabidopsis to be essential for catalytic activity as guanine nucleotide exchange factor of ROP (Gu *et al.,* 2006; Oda *et al.,* 2012). (C) Unrooted Maximum-Likelihood tree of KAR and the other related RopGEF proteins across various plant lineages. The numbers on the branches show bootstrap values calculated from 1000 replicates. The scale bars are evolutionary distance at the ratio of amino acid substitutions.

**Figure S4.**
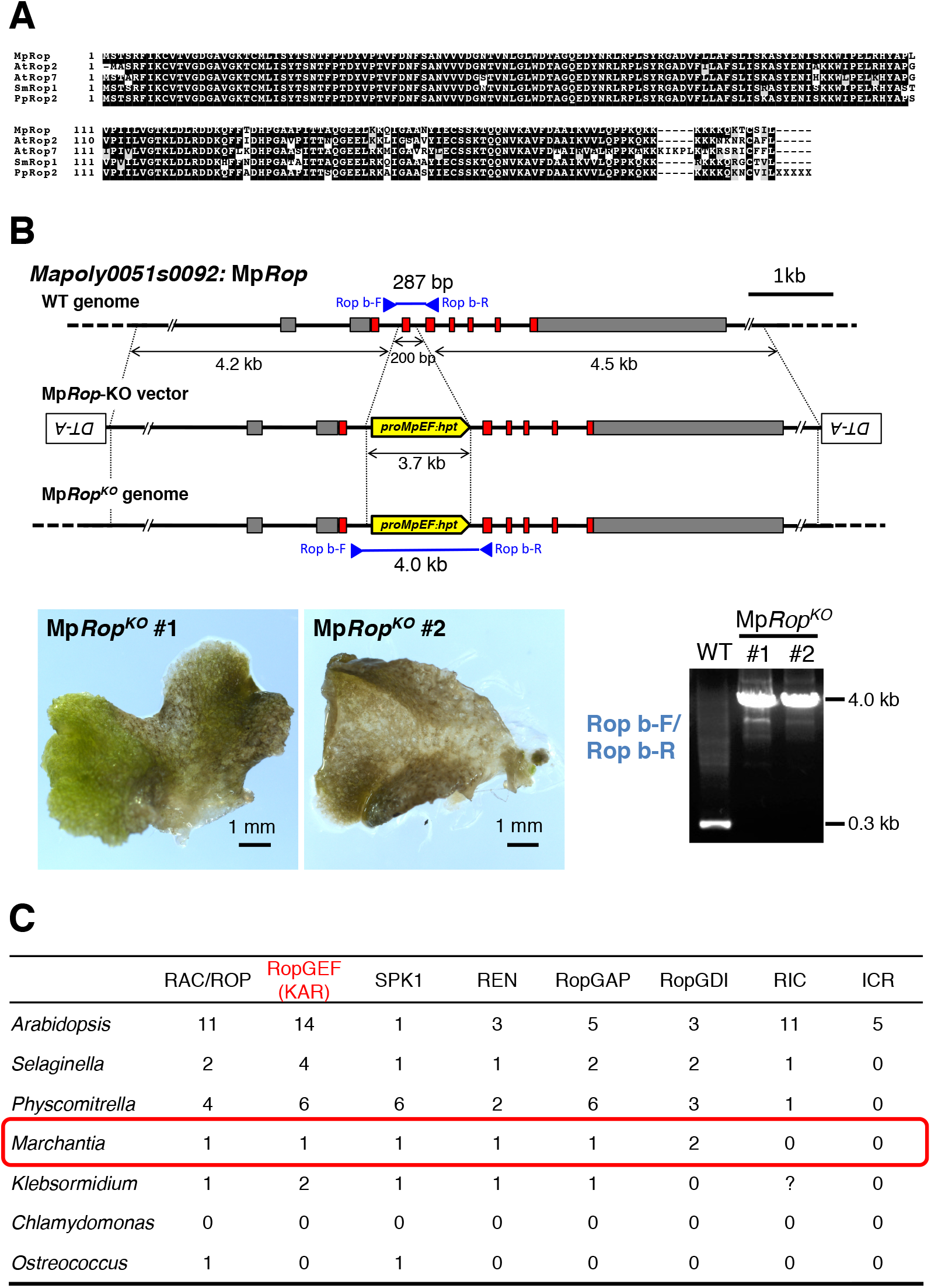
**RopGTPase and its related gene families in *M. polymorpha*, Related to Figure 4** (A) Multiple alignment of the full-amino acid sequences of MpROP and representative RopGTPases in *P. patens*, *S. moellendorfii*, and *A. thaliana*. (B) Generation of knockout mutants of Mp*Rop*. Schematic representation of the structure of the Mp*Rop* locus in wild type, the construct designed for gene targeting, and the Mp*Rop* locus disrupted in the gene-targeted lines (upper). Phenotype of two independent disruptants of Mp*Rop* gene showed severe impairment of thallus growth, and had died in the early stage of thallus development (lower left). A genotyping of Mp*Rop* indicates successful disruption of the cds structure occurred in Mp*Rop* knockouts (lower right). (C) Relative sizes of Rop signaling gene families in Viridiplantae. Total number of all homologous genes in the indicated gene families are indicated.

**Table S1:**
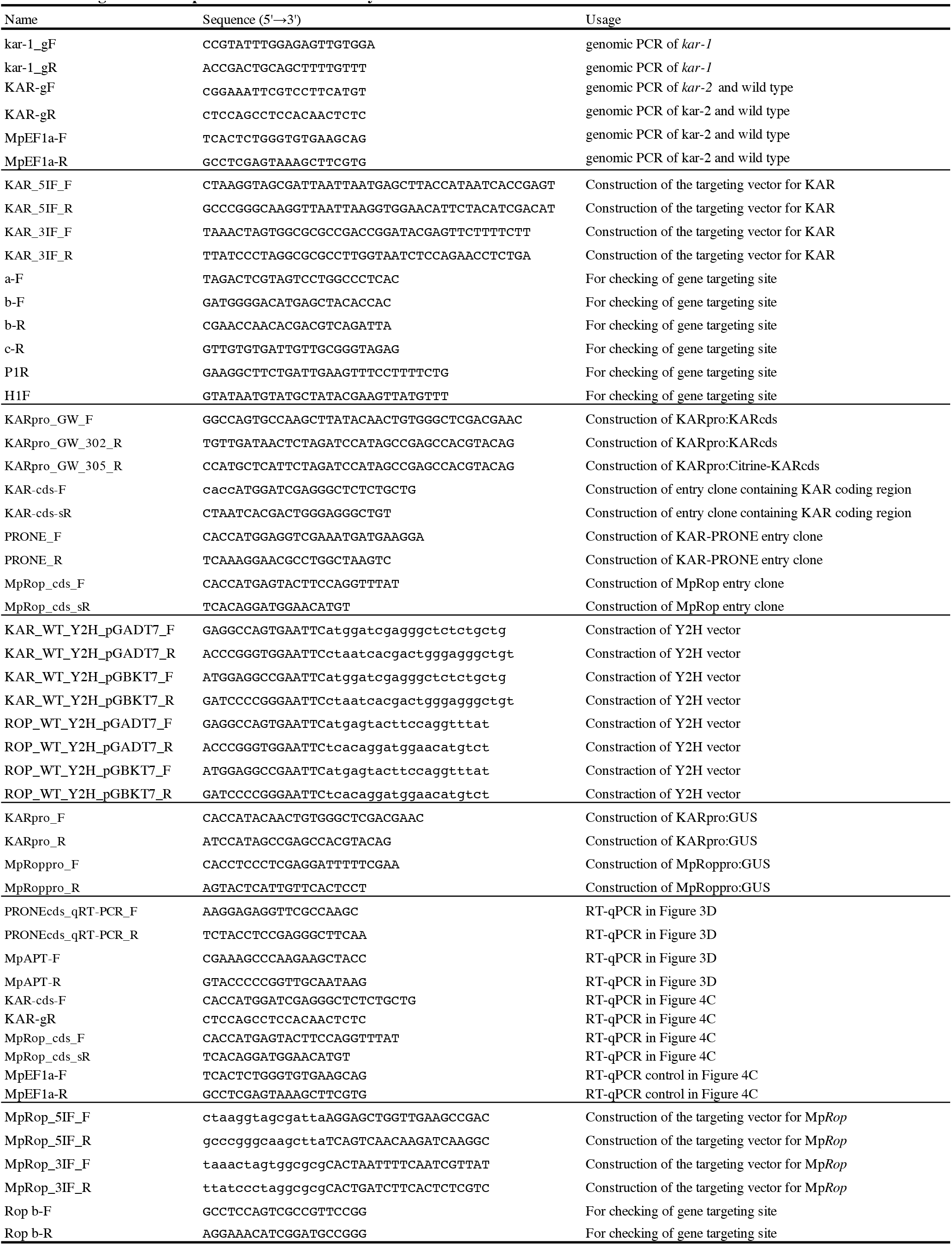
Oligonucleotide primers used in this study.

